# Incorporation of DNA methylation into eQTL mapping in African Americans

**DOI:** 10.1101/2020.08.05.238030

**Authors:** Anmol Singh, Yizhen Zhong, Layan Nahlawi, C. Sehwan Park, Tanima De, Cristina Alarcon, Minoli A. Perera

## Abstract

Epigenetics is a reversible molecular mechanism that plays a critical role in many developmental, adaptive, and disease processes. DNA methylation has been shown to regulate gene expression and the advent of high throughput technologies has made genome-wide DNA methylation analysis possible. We investigated the effect of DNA methylation in eQTL mapping (methylation-adjusted eQTLs), by incorporating DNA methylation as a SNP-based covariate in eQTL mapping in African American derived hepatocytes. We found that the addition of DNA methylation uncovered new eQTLs and eGenes. Previously discovered eQTLs were significantly altered by the addition of DNA methylation data suggesting that methylation may modulate the association of SNPs to gene expression. We found that methylation-adjusted eQTLs which were less significant compared to PC-adjusted eQTLs were enriched in lipoprotein measurements (FDR = 0.0040), immune system disorders (FDR = 0.0042), and liver enzyme measurements (FDR = 0.047), suggesting a role of DNA methylation in regulating the genetic basis of these phenotypes. Our methylation-adjusted eQTL analysis also uncovered novel SNP-gene pairs. For example, our study found the SNP, rs11546996, was associated to *PNKP.* In a previous GWAS, this SNP was associated with primary biliary cirrhosis although the causal gene was thought to be *SPIB*. Our methylation-adjusted method potentially adds new understanding to the genetic basis of complex diseases that disproportionally affect African Americans.

## 1. Introduction

DNA methylation plays an important role in the regulation of gene expression which in turn affects many complex diseases and traits.^1^ However, integrating multi-omics data, such as DNA methylation and expression Quantitative Trait Loci (eQTL) mapping, can be challenging as the addition of SNP-based covariates is computationally intensive and multi-omics datasets with matching samples are sparse.^2^ Moreover, matching datasets in minority populations are nearly absent from public databases. DNA methylation patterns, in particular, are very complex, vary greatly from sample to sample^3^, and change due to environmental factors.^4^ Therefore, DNA methylation studies are hard to generalize. The advent of high throughput and next generation sequencing technologies, however, has made it possible for DNA methylation to be analyzed genome-wide.^4^ Several investigators have previously integrated genome-wide sequencing data and DNA methylation to uncover SNPs that significantly associate to CpG methylation, called methylation QTLs (meQTLs).^5-7^ These studies have found that methylation plays a significant role in the onset of diseases and phenotypes such as obsessive-compulsive disorder, prostate cancer, and drug response.^5-7^ Most of these studies have been conducted in populations of European ancestry.

The African American population is widely underrepresented in genetic studies which is shown by the distribution of individuals participating in GWAS studies. In GWAS studies, only 19% of individuals are non-European and less than 5% are non-European and non-Asian.^8^ While other eQTL mapping studies have used African American samples, the number of individuals have been very small, thus making them underpowered to adequately account for population specific variation. Furthermore, these studies did not account for methylation as a SNP-based covariate.^8^ In this study, we perform the first investigation of the effect of DNA methylation on eQTL mapping in African Americans and how eQTL associations to complex diseases, phenotypic traits, and metabolic traits may be modulated by DNA methylation. These findings may help explain the role DNA methylation plays in health disparities observed in African Americans.

## 2. Methods

### 2.1. Cohort

Sixty-eight African American hepatocyte cell cultures were acquired. After genotyping, DNA methylation quality control and RNA-sequencing quality control, 53 samples were used to conduct this study as shown in Fig. 1. Hepatocytes were either purchased from commercial companies (BioIVT, TRL, Life technologies, Corning, and Xenotech) or isolated from cadaveric livers using the same procedure described in Park et. al.^9^ All genomic, transcriptomic and methylome data were gathered from the same hepatocyte samples.

**Fig. 1.**
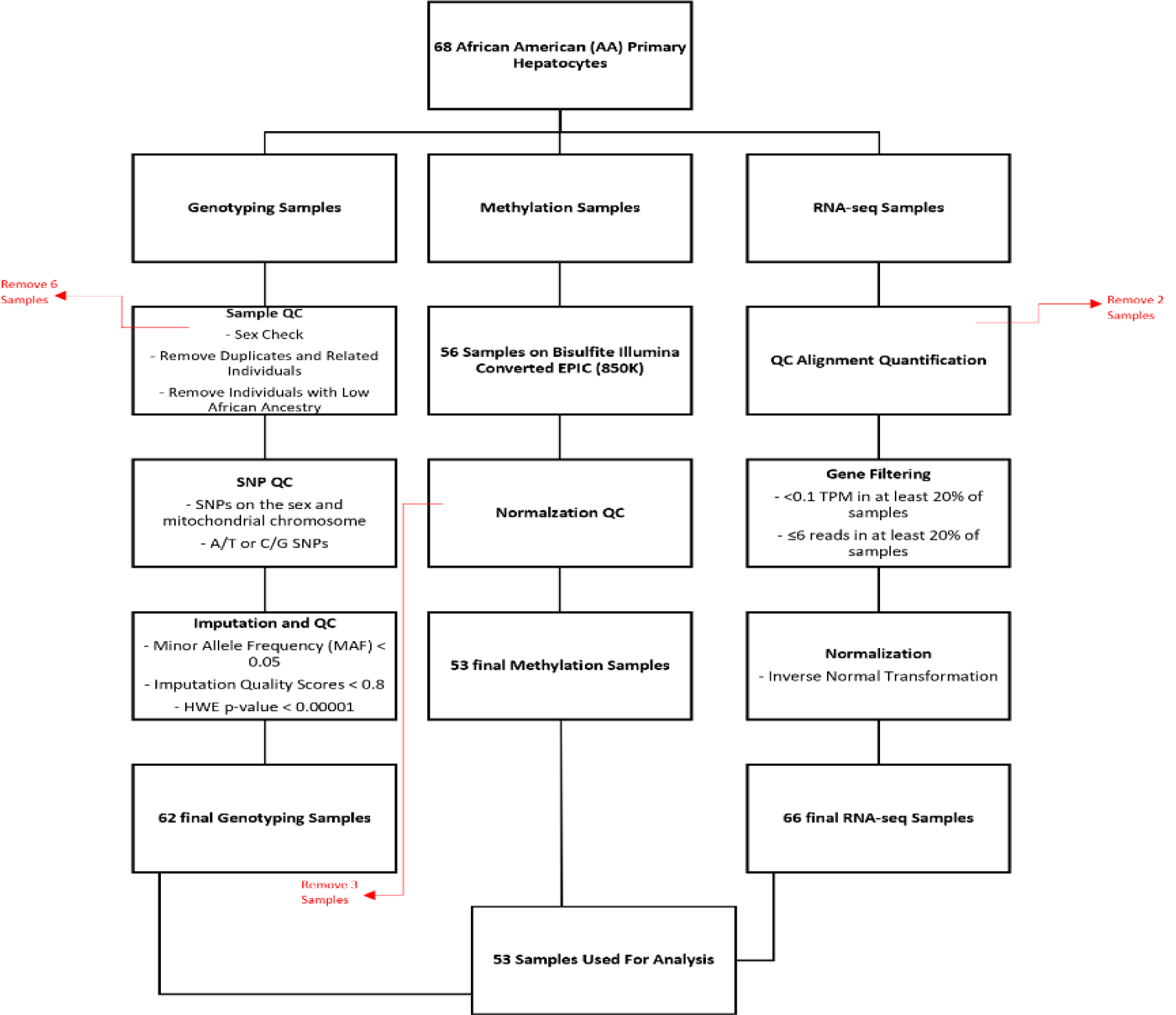
Flowchart showing the study design and the methods used in each dataset.

### 2.2. Genotyping, Imputation, and QC

DNA was extracted from each hepatocyte culture using Gentra Puregene Blood kit (Qiagen) and all the DNA samples were bar coded for genotyping. The SNPs were then genotyped using the Illumina Multi-Ethnic Genotyping array (MEGA) at the University of Chicago Functional Genomics Core using standard protocols. The outputs were then created by Genome Studio using a 0.15 GenCall score as the cutoff. PLINK^10^ was then used to perform a sex check and to identify individuals with contradictory sex information. The identity-by-descent method was used with a cut off score of 0.125 to identify duplicated or related individuals, where the cutoff score indicates third-degree relatedness. Furthermore, samples that did not cluster with African American (AA) samples on the PCA plot were removed. This left a total of 62 samples, with 6 samples being removed during these steps. SNPs were excluded from the analysis based on the criteria described in Fig. 1 under SNP QC and Imputation and QC as previously described in Park et. al.^9^, leaving 7,180,502 SNPs used in the analysis.

### 2.3. RNA-sequencing and QC

Total RNA was extracted from each primary cell culture after three days of plating using the Qiagen RNeasy Plus mini-kit. Only the samples with RNA integrity number (RIN) score > 8 were sequenced. RNA-seq libraries were prepared using TruSeq RNA Sample Prep Kit, Set A (Illumina catalog # FC-122-1001) in accordance with the manufacturer’s instructions. Illumina HiSeq 2500 and HiSeq 4000 machines were used to prepare the cDNA libraries sequence and. This resulted in 50 million reads per sample (single-end 50bp reads).

Quality of the raw reads from FASTQ files was assessed by FastQC (v0.11.2). A per base sequence quality threshold of > 20 across all bases was set for the fastq files. STAR 2.5^11^ was used to align the reads to human Genome sequence GRCh38 and Comprehensive gene annotation (GENCODE version 25). Only uniquely mapped reads were retained and indexed by SAMTools 1.2.^12^ To assess the nucleotide composition bias, GC content distribution and coverage skewness of the mapped reads read_NVC.py, read_GC.py and geneBody_coverage.py from RNA-SeQC (2.6.4) were used. Lastly, Picard CollectRnaSeqMetrics was applied to evaluate the distribution of bases within transcripts. Fractions of nucleotides within specific genomic regions were measured and only samples with > 80% of bases aligned to exons and UTRs regions were retained for analysis.

### 2.4. Gene expression quantification

To quantify gene expression a collapsed gene model was used, following the GTEx isoform collapsing procedure.^13^ The reads were mapped to genes referenced with Comprehensive gene annotation (GENCODE version 25) to evaluate gene-level expression using RNA-SeQC.^14^ The Bioconductor package, DESeq2 (version1.20.0)^15^ was used to supply HTSeq^16^ raw counts for the analysis of gene expression. DESeq2 was also used to perform principal component analysis (PCA). Using regularized log transformation, the counts were normalized. PC1 and PC2 were plotted to visualize the expression patterns of the samples and two samples with distinct expression patterns were excluded as outliers.

The gene expression was normalized by the trimmed mean of M-values normalization method (TMM), which was implemented in edgeR.^17^ The TPM (transcript per million) was calculated by first normalizing the counts by gene length and then normalizing by read depth. The thresholds for gene expression values were set at < 0.1 TPM in at least 20% of samples and ≤ 6 reads in at least 20% of samples. Inverse normal transformation was used to normalize the expression values for each gene. The gene coordinates were remapped to hg19/GRCh 37 (GENCODE version 19).

### 2.5. Methylation Sample Preparation and QC

DNA was isolated from hepatocytes as described in Park et al.^9^ As shown in Fig. 1, only 56 of the hepatocyte samples produced sufficient bisulfite-converted DNA for analysis. The Illumina MethylationEPIC BeadChip microarray (San Diego, Ca, USA), consisting of approximately 850,000 probes^18^ was used for methylation profiling of DNA extracted from 56 AA hepatocytes that overlapped the samples used for gene expression analysis.

Methylation data QC and normalization was performed using the ChAMP R package (version 2.10.1)^19^ as previously described in Park et.al.^9^ This process removed: 9204 probes for any sample that did not have a detection p value <0.01 which is counted as a failed probe, 1043 probes with a bead count <3 in at least 5% of samples, 49 probes that align to multiple locations as identified by Nordlund et al.^20^, 2975 probes with no CG start sites, and 17,235 probes located on X and Y chromosomes. From the QC process, three samples were excluded as outliers resulting in 53 samples remaining in the analysis.

### 2.6. Methylation-adjusted eQTLs

The R package Matrix eQTL^21^ was used to determine the methylation site(s) that correspond to each SNP within a 2.5 kB window. The resulting file was then analyzed to check if the identified CpG sites were within 2.5 kB of the corresponding SNP. CpG sites were then grouped together by SNP to determine the number of CpG sites on average at each SNP and to determine the pairwise correlation between CpG sites at each SNP. We used a weighted average based on the distance of the CpG site from the SNP to determine the methylation values for each SNP. If only one CpG site was linked to a SNP, then the weight of the CpG site would be:

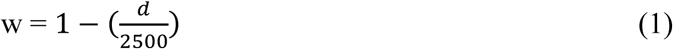

where d is the genomic distance (in base pairs) between the CpG site and the SNP and 2500 represents the 2.5kB window size used in this analysis. This weight would then be multiplied by the methylation value of the CpG site to get the normalized methylation value used in the analysis. This weighting system allowed proximal CpG sites to have a greater weight. If more than one CpG site was found within the 2.5kB region then each CpG site’s weight, *w*i, was calculated using equation (1) above and the final weight for each CpG site was calculated as:

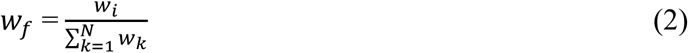

where N is the total number of CpG sites that correspond to a particular SNP and 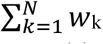 represents the sum of the initial weights of all the CpG sites that correspond to that SNP. This calculation ensures the sum of the final weights of all CpG sites corresponding to a single SNP are equal to one. The SNP-based methylation value was then calculated by:

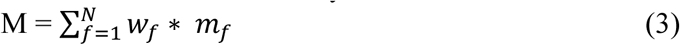

where M is the SNP-based methylation value and 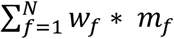 represents the weighted sum of all of the methylation values for the CpG sites corresponding to that SNP. These averaged methylation values used as a SNP based covariate and eQTLs were mapped using the LAMatrix R package.^22^ The methylation-adjusted eQTLs were then compared with the PC-adjusted eQTLs (mapped in the same hepatocyte cultures) to investigate if there were any changes in the significance of the previously published eQTL findings.^22^

### 2.7. eQTL and GWAS overlap

To understand how the methylation-adjusted eQTLs may explain the underlying mechanisms in GWAS findings, the method presented in Zhong et. al.^8^ was used with some modifications. This included downloading the NHGRI/EBI GWAS Catalog file (v.1.0.2, 2019-03-22) and keeping only the associations that passed the genome-wide significant level (p<5e-8). Furthermore, the rsids were remapped from Build38 to Build37 using Ensembl API. The 1000 Genomes YRI population were used to extract all the variants in LD with the independent GWAS variants (r^2^ > 0.8) and the traits of the corresponding GWAS hits were put into 17 groups which corresponded to ontology-based trait categories.^23^ A false discovery rate (FDR) threshold of 0.05 was set as significant enrichment for an ontology. The methylation-adjusted eQTLs were split into three groups for this analysis: (i) eQTLs that were significant with PC-adjustment and increased in significance with methylation-adjustment, (ii) eQTLs that were not significant with PC-adjustment and became significant with methylation-adjustment (FDR<0.05), and (iii) eQTLs that were significant with PC-adjustment and became less significant with methylation-adjustment. These three groups of eQTLs were compared to the GWAS variants.

## 3. Results

Fifty-three African American hepatocyte samples were used in this analysis, with 28 (52.8%) males and 25 (47.2%) females. The age (mean ± standard deviation) of the cohort was 39 ± 20.5 years old. To account for methylation in this eQTL mapping analysis, the LAMatrix method was used.^22^ Instead of incorporating local ancestry into the analysis as previously done^8^, DNA methylation was used in its place. LAMatrix was chosen because the R package has increased power and controls false positives when gene expression differs by locus-specific covariate, such as methylation.^22^ The genotype-gene expression associations within a *cis* region (1 Mb around the gene) were tested. Sex, platform, batch, and 10 PEER variables estimated from normalized expression values were used as covariates in the analysis as previously described in Zhong et.al.^8^ We chose the number of PEER variables to use in our analysis in relation to the same study by Zhong et. al.^8^

### 3.1 Methylation-adjusted *eQTLs vs PC-adjusted eQTLs*

Out of the 7,180,502 total SNPs in the dataset, 2,494,181 SNPs had at least one CpG site within the kB window, with an average of 3.08 CpG sites per SNP (ranging between of 1 to 95 CpG sites per SNP). We identified 2,296 eQTLs with methylation-adjustment at an FDR threshold of 0.05. To ascertain if any methylation-adjusted eQTLs resulted in the novel discovery of regulatory variation, we compared significant methylation-adjusted eQTLs (FDR<0.05) against significant PC-adjusted eQTLs (FDR<0.05). This comparison resulted in 308 unique methylation-adjusted eQTLs that were not found with PC-adjusted analysis, and 1,954 eQTLs which were common to both analyses. The remaining 19,567 found in PC-adjustment were not significant in this analysis (Fig. 2A). We compared the effect size for those eQTLs found in both analyses and they did not differ significantly.

**Fig. 2.**
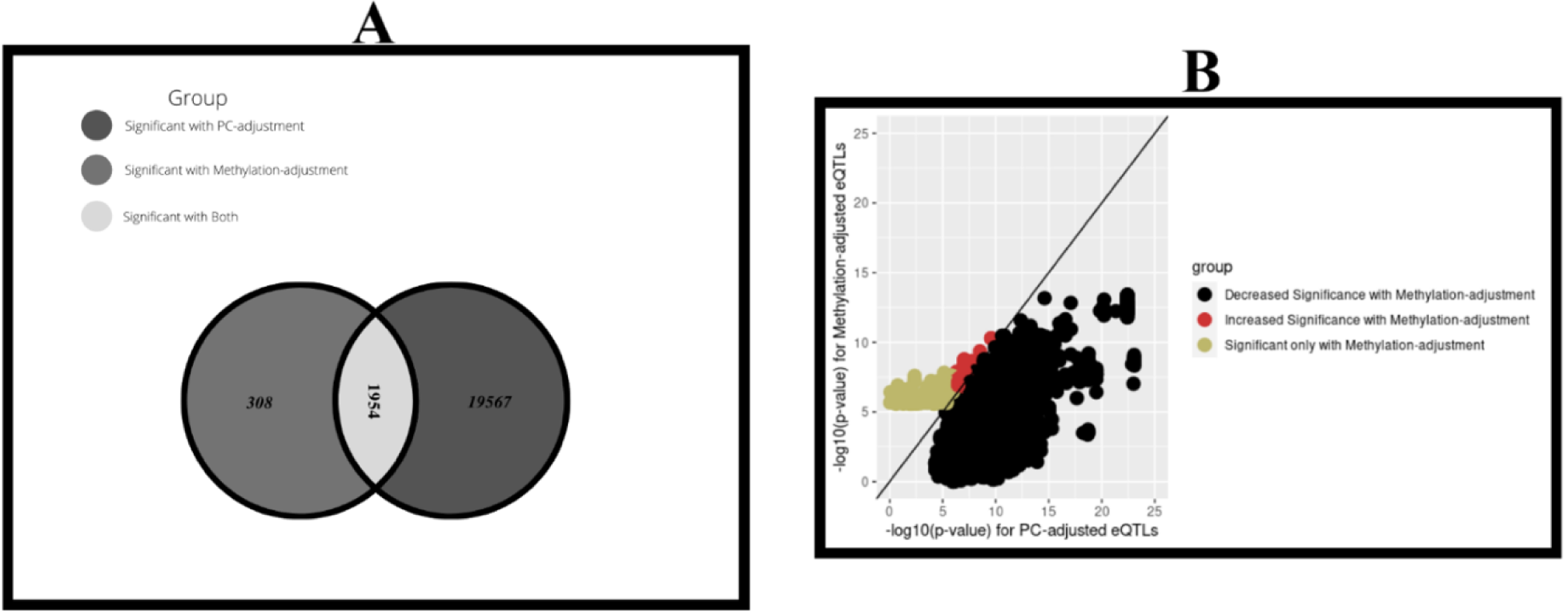
Methylation-adjusted eQTLs as compared to PC-adjusted eQTLs. A) The Venn-Diagram shows the number of eQTLs that are significant only with Methylation-adjustment, significant only with PC-adjustment, and those that are significant in both analyses. B) This plot compares the p-values of the 308 eQTLs that are significant only with methylation-adjustment, 11,485 that were significant with PC-adjustment and decreased in significance with methylation-adjustment, and 50 that were significant with PC-adjustment and increased in significance with methylation-adjustment. There are also 9,679 eQTLs that showed no change in p-value as they did not have nearby methylation sites or methylation had no effect on the association.

Finally, the comparison revealed that there were 11,485 eQTLs that were significant with PC-adjustment and decreased in significance with methylation-adjustment and 50 eQTLs that were significant with PC-adjustment increased in significance with methylation-adjustment (Fig. 2B).

### 3.2 GWAS Associations for Methylation-adjusted eQTLs

We verified the role of these methylation-adjusted eQTLs in the pathogenesis of complex diseases or traits by overlapping the methylation-adjusted eQTLs with SNPs in previously reported GWAS. For this analysis, variants, from NHGRI-EBI GWAS catalog, or their tagging variants (r^2^ > 0.8, 1000 Genomes YRI population) were used to determine which GWAS SNPs intersect with the methylation-adjusted eQTLs. To analyze the effect of methylation even further, the methylation-adjusted eQTLs were broken into three groups: (i) eQTLs that are only significant with methylation-adjustment, (ii) eQTLs that were significant with PC-adjustment but became more significant with methylation-adjustment, and (iii) eQTLs that were significant with PC-adjustment but became less significant with methylation-adjustment. In total there were 285 GWAS associations that intersect with methylation-adjusted eQTLs across the three groups.

#### 3.2.1. Group 1: eQTLs that were only significant with methylation-adjustment

For eQTLs that were only significant with methylation-adjustment, 16 GWAS associations were found that intersected with these eQTLs. There was significant enrichment for digestive system disorders (FDR = 0.011), as shown in Fig. 3A. One of the eQTLs enriched for digestive system disorders, rs11546996, was associated with primary biliary cirrhosis as shown by Hirschfield et. al.^24^ Due to the intergenic location of rs11546996 within *SPIB*, the causal gene was reported as *SPIB* in this study. However, our analysis associated rs11546996 to *PNKP* (P-value = 1.05e-6, FDR = 0.026) thereby potentially identifying a new causal gene for primary biliary cirrhosis by accounting for methylation in eQTL mapping.

**Fig. 3.**
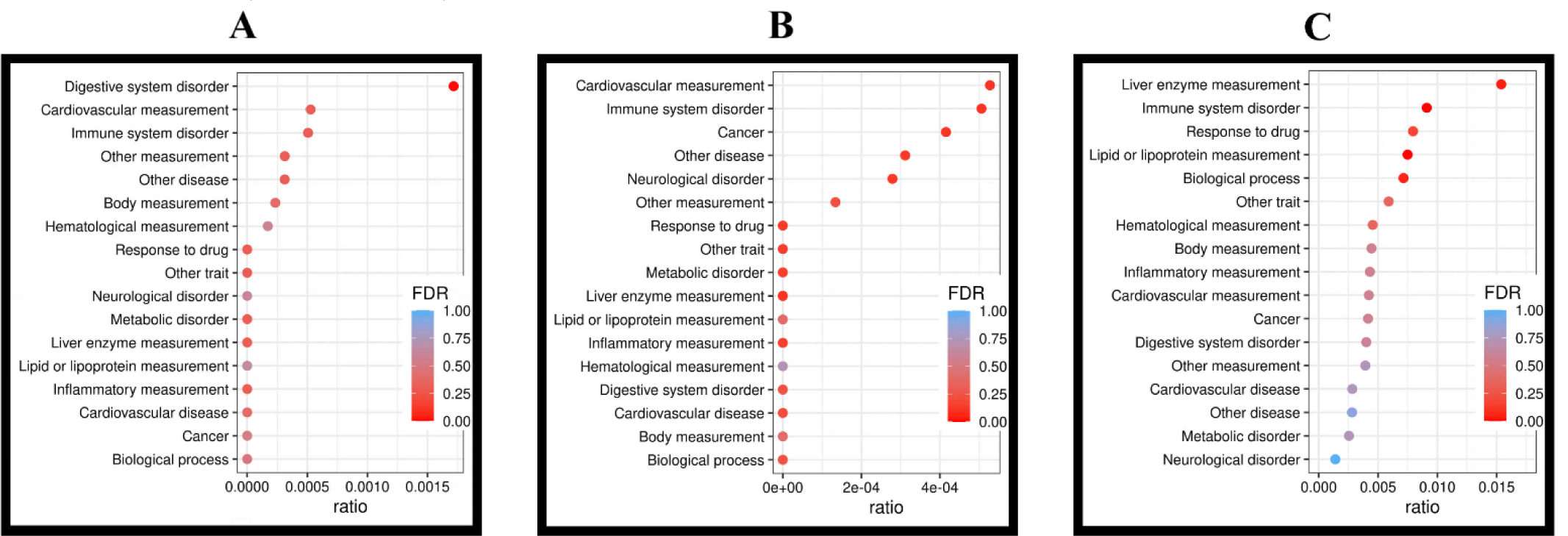
Enrichment of methylation-adjusted eQTLs in GWAS findings. These figures show the enrichment in GWAS categories for different sets of methylation-adjusted eQTLs: A) eQTLs, that were only significant with methylation-adjustment. B) eQTLs, that were significant with PC-adjustment and increased in significance with methylation-adjustment. C) eQTLs, that were significant with PC-adjustment and decreased in significance with methylation-adjustment. In each figure the dot represents the proportion of SNPs within each category that were also within the eQTL group on the x-axis and FDR for enrichment is shown by the color of the dot.

#### 3.2.2. Group 2: eQTLs that were significant with PC-adjustment and increased in significance with methylation-adjustment

For eQTLs that were significant with PC-adjustment and increased in significance with methylation-adjustment, eight GWAS associations intersected with this group of eQTLs. None of them were significantly enriched for any ontology (FDR<0.05), as shown in Fig. 3B. This could be due to the very small sample size of eQTLs in this group.

#### 3.2.3. Group 3: eQTLs that were significant with PC-adjustment and decreased in significance with methylation-adjustment

For eQTLs that were significant with PC-adjustment and decreased in significance with methylation-adjustment, 261 GWAS associations intersected with eQTLs in this group. There was significant enrichment for lipid or lipoprotein measurements (FDR = 0.0040), immune system disorders (FDR = 0.0042), and liver enzyme measurements (FDR = 0.047), as shown in Fig. 3C. This suggests a more complex interplay between SNPs and genes implicated in these diseases, such that these SNPs may be associated to susceptibility of disease, but that susceptibility may be modulated by DNA methylation.

Novel SNP-gene associations were also found. Two examples are rs7528419 and rs12740374, which are associated with *SORT1*, a gene known to influence LDL-cholesterol levels and lipid/lipoprotein measurements.^25, 26^ When accounting for methylation the p-value of these two SNP-gene pairs increased from 1.99e-9 for both to 4.92e-8 and 1.25e-7, respectively. The FDR also increased from 3.01e-5 for both to 3.12e-3 and 6.08e-3, respectively. Furthermore, both SNPs had a low amount of methylation, ranging from 0.12 to 0.33. Although these SNP-gene pairs remained significant with methylation-adjustment (FDR<0.05), they’re significance decreased dramatically indicating that methylation, between 0.12 and 0.33 near these SNPs, may play a role in the association between these SNPs to lipid phenotypes. This suggests that DNA methylation should be considered when assessing genomic risk of LDL-cholesterol levels and cholesterol-related diseases, such as myocardial infarction.

Another methylation-adjusted eQTL, rs9296736 associated with the expression of *MLIP*, was previously found to be associated with liver enzyme measurements.^27^ High levels of liver-enzymes in plasma are widely associated with an increased risk for developing many diseases including cirrhosis and cardiovascular disease.^27^ This SNP-gene pair decreased in significance considerably when it was adjusted for methylation. The p-value and FDR of this methylation-adjusted eQTL went from 2.08e-9 and 3.13e-5 with PC-adjustment to 0.037 and 0.94, respectively, with methylation-adjustment. For this SNP-gene pair, rs9296736 was highly methylated, with methylation values ranging from 0.89 to 0.97. This result suggests that the association of rs9296736 to *MLIP* and liver enzyme measurements may depend on the surrounding DNA methylation landscape.

### 3.3. Discovery of eGenes associated to disease traits using methylation-adjusted eQTL mapping

There were 179 eGenes found through methylation-adjusted eQTL mapping (FDR<0.05) as well as 80 eGenes that were not significant with PC-adjustment. Two of these eGenes, *GSTM3* (FDR = 0.014) and *HSPA6* (FDR = 0.029), have been associated to disease traits such as Hepatitis B (HBV) for *GSTM3* and Hepatocellular Carcinoma (HCC) for both eGenes.^28-31^ African Americans have a higher incidence and worse outcomes of HBV and HCC when compared to other demographics.^32, 33^ Since these eGenes were not significant with PC-adjusted eQTL mapping, they may explain how methylation plays a role in the health disparities observed in African Americans. As shown in Fig. 4, there is a significant correlation between rs1332018 genotype and *GSTM3* expression as well as rs1332018 genotype to DNA methylation. From this we can see that the T allele is associated with both great gene expression and low DNA methylation. A total of 18 CpG sites contributed to this association. As shown in Fig. 5, there is also a direct relationship between *HSPA6* gene expression and DNA methylation with the A allele associated with both greater gene expression and increased DNA methylation. A total of 7 CpG sites contributed to this association.

**Fig. 4.**
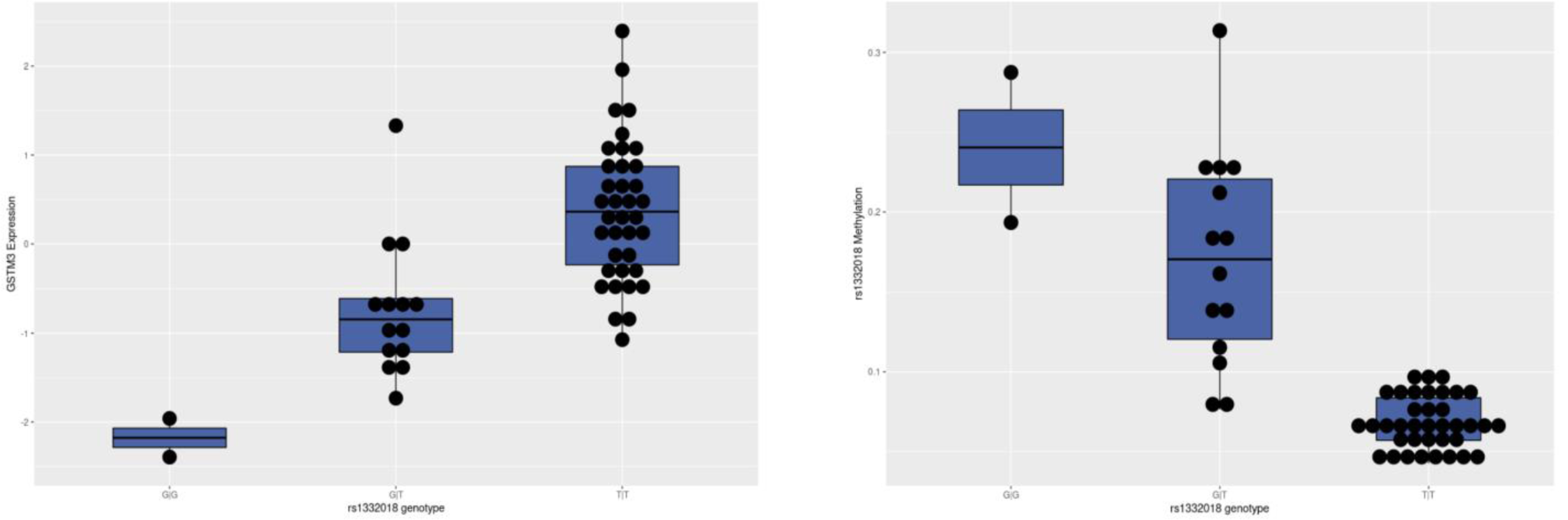
Boxplots of rs1332018 genotype vs *GSTM3* expression and rs1332018 methylation. This figure shows the significant increase in *GSTM3* gene expression with the leads to a significant decrease in methylation.

**Fig. 5.**
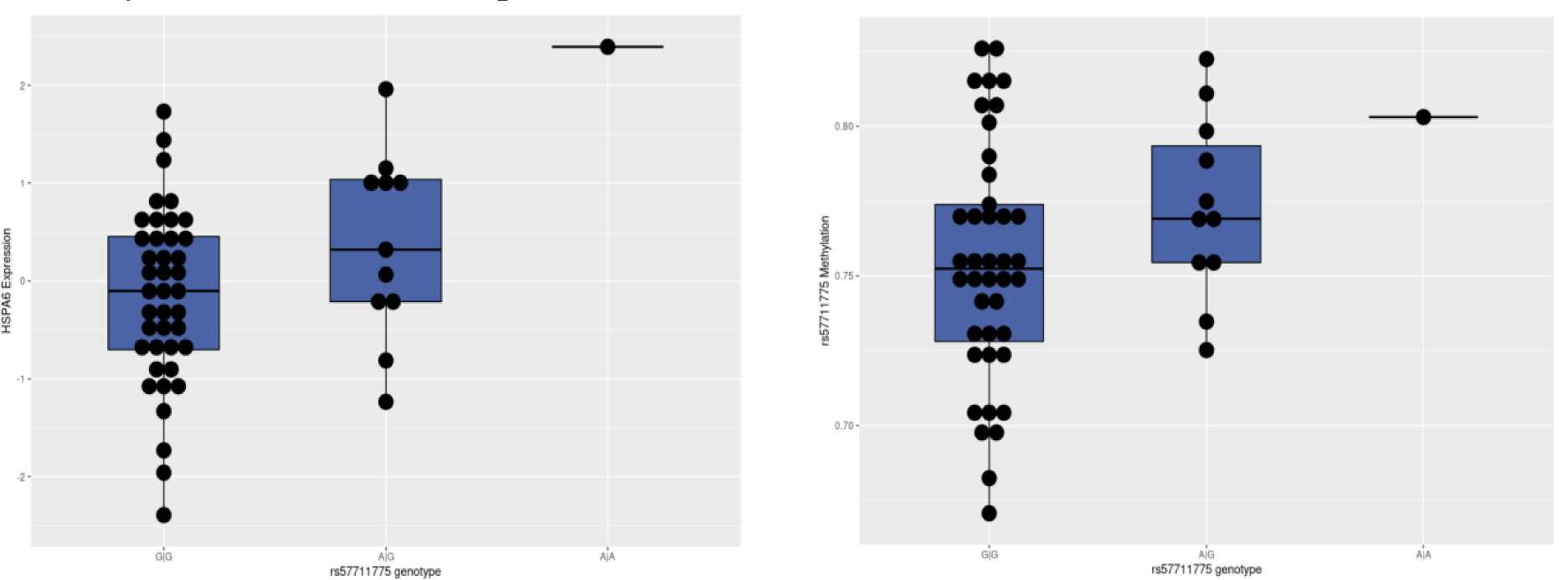
Boxplots of rs57711775 genotype vs *HSPA6* expression and rs57711775 methylation. This figure shows that significant increase in *HSPA6* expression leads to a significant increase in rs57711775 methylation

## 4. Discussion

Through the integration of DNA methylation into eQTL mapping, we showed how methylation potentially plays a critical role in SNP-gene associations as well as the association of these eQTLs to diseases and metabolic traits. Our analysis was aided by the use of the computationally efficient R package, LAMatrix, which allows for the addition of a SNP based covariate to eQTL mapping. Additionally, our use of data from African Americans aided in the discovery of new regulatory variants as this population is more genetically diverse than European ancestry populations.

We found unique eGenes in our analysis that were not found by eQTL mapping with only PC-adjustment. Two of these eGenes, *GSTM3* and *HSPA6*, are associated with diseases such as HBV for *GSTM3* and HCC for both eGenes.^28-31^ These are diseases that disproportionately affect African Americans.^32, 33^ *GSTM3* has also been associated to oxidative stress and specifically several studies have found that epigenetic suppression of *GSTM3* in HBV-infected cells causes oxidative stress^29, 30^, which can lead to HCC.^28^ Furthermore, other studies showed that *GSTM3* expression was lowered with promoter hypermethylation^34^ and in chemical-induced HCC.^35^ This agrees with the previous studies mentioned, showing that epigenetic suppression of *GSTM3* leads to HCC in HBV-infected cells.^28-30^ We found a significant inverse relationship between *GSTM3* expression and DNA methylation around rs1332018. This suggests that individuals with rs1332018 genotypes that have a lower *GSTM3* expression and higher methylation may be at a higher risk for HCC. *HSPA6* was also found to be overexpressed in human HCC tissues and a potential risk factor for HCC reccurence.^31^ We found that the expression of *HSPA6* increased with methylation around rs57711775, which could mean that methylation potentially plays a role in upregulating *HSPA6*. Furthermore, the A allele of rs57711775 that is associated with higher *HSPA6* expression and higher methylation in our analysis is not found at all in European ancestry populations according to the Ensembl database. Thus, we potentially elucidated a causal variant and risk allele for HCC specific to African Americans by using methylation in this analysis. As has previously been reported, the direction of effect of DNA methylation is dependent on the location of methylation.^36, 37^ Previous studies have shown that methylation within the transcriptional start site of the promoter is well known to repress gene expression while methylation within the gene body results in more variable expression.^36, 37^ Therefore, both *GSTM3* expression and *HSPA6* expression may contribute to the onset of HCC in African Americans.

In our GWAS enrichment, we found a significant enrichment for digestive system disorders for the eQTLs significant only with methylation-adjustment and a significant enrichment for lipid or lipoprotein measurements, immune system disorders, and liver enzyme measurements for the eQTLs that are less significant with methylation-adjustment. Both immune-related phenotypes and lipid and lipoprotein measures differ by population and may contribute to disease disparities. Our findings suggest that methylation may play a role in these diseases. Further studies are needed to determine if methylation at these specific SNPs and genes differ between populations. This analysis also revealed an interesting association with eQTLs that were only significant after methylation-adjustment. The SNP, rs11546996, a SNP associated with primary biliary cirrhosis, was a methylation-adjusted eQTL for *PNKP*. In a previous GWAS study, a causal SNP-gene association for primary biliary cirrhosis was found with rs11546996 and the causal gene was assumed to be *SPIB*, as it is the closest gene.^24^ Since our study specifically looked at gene expression in hepatocytes, a tissue relevant for this disease, we may have found a potentially novel SNP-gene pair associated with primary biliary cirrhosis whose expression is regulated by both methylation and gene variation. *PNKP* has also been associated with repairing DNA after damage from oxidative stress^38^, so rs11546996 could be a SNP that aids in this process.

There were several limitations to our study. First, we were only able to include 53 samples into this analysis and hence our analysis was underpowered. Second, we assessed methylation with the Illumina EPIC array which is limited to the CpG sites chosen for the chip. Unmeasured DNA methylation may have effects on eQTLs that were not captured by our analysis. Third, our results, compared to the findings in the entire GWAS catalog, are only applicable to diseases in which hepatocytes play a key role. Our findings may not be generalizable to other cell or tissue types. Finally, we have assumed that DNA methylation closer to the SNP is more likely to influence eQTL mapping, however this may not always be the case. With greater meQTL analysis in relevant cell types and populations, we may be able to weight the effect of methylation more precisely as a SNP-based covariate.

In conclusion, this is the first study to explore the effect of DNA methylation in eQTL mapping in African Americans. The African American demographic is widely underrepresented in genetic studies and their greater genetic diversity may allow us to find novel SNP-gene pairs as well as population specific SNPs. Our findings can be used to understand how DNA methylation potentially plays a role in complex diseases, phenotypic traits, and metabolic traits in African Americans.

## 5. Acknowledgments

This work was supported by NIH National Institute on Minority Health and Health Disparities (NIMHD) Research Project 1R01MD009217-01 (R01).

